# Sequential visual stimuli increase high frequency power in the visual cortex

**DOI:** 10.1101/2025.01.14.632930

**Authors:** Julian Keil, Victor Hernandez-Urbina, Chrystalleni Vassiliou, Camin Dean, Dietmar Schmitz, Jens Kremkow, Jérémie Sibille

## Abstract

Today, 40 Hz flickering full-field visual stimulation is used to entrain neuronal oscillations for a variety of therapeutic purposes. We here propose spatially organized sequential visual flickering stimulation as a newer tool to entrain the visual system. We show that sequential visual flickering can evoke increased power in high frequencies (100 to 190 Hz) in the visual cortex of mice. Consequently, sequential sensory stimulation should be regarded as a putative new way leading to power increases in high frequency domains.

## MAIN TEXT

Visual neuronal entrainment with 40 Hz full-field stimulation has been recently established as a neuroprotective approach to counteract neurodegeneration associated with Alzheimer’s disease^1,2^, slow down the disease progression^3^, and improve sleep^4^. However, the actual benefit of visual full-field 40 Hz gamma entrainment against neurodegeneration has been recently questioned^5^. These concerns are further strengthened by an independent study on the intrinsic properties of the functional transmission of full-field visual stimulation *in vivo*. The authors report a progressive attenuation of the desired 40 Hz neuronal entrainment along the visual pathway, leading to an absence of 40 Hz entrainment in the hippocampus, probably in part due to biophysical neuronal properties^6^. Overall, neuronal tissue tends to dissipate oscillation power, acting as a low pass filter with a 1/frequency characteristic^7^. Thus, the few instances where the local field potential (LFP) is transiently increased are of particular interest. To overcome such low pass limitations, we propose to spatially combine neighboring flickering checkerboard patterns with very small delays along different retinotopic positions^8^ (Fig. 1C). These sequential visual flickering stimulations take advantage of the independence of parallel retinotopic pathways in order to activate different positions with very short delays in order to stimulate the cortex in a predetermined way.

**Figure 1:**
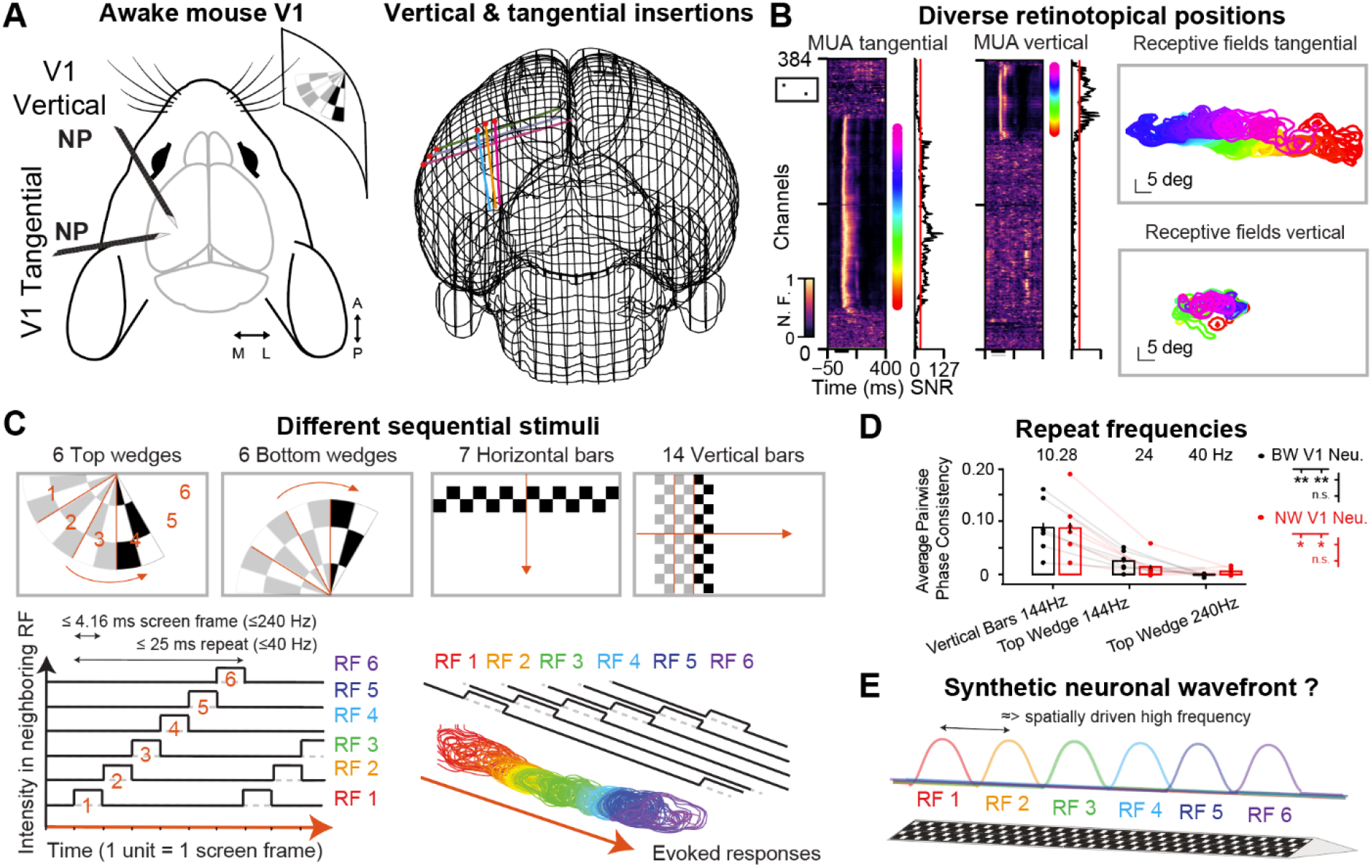
Neuropixels probe insertions to measure sequential wavefront propagation throughout the retinotopy. **A**. Schematic of a mouse passively viewing stimulations presented on a screen, with tangential and vertical Neuropixels (NP) insertions (left), and the 3D reconstruction of the brain showing probe placements (right). **B**. Visually driven multi-unit-activity (MUA) evoked by black squares on white background for tangential (left) and vertical (right) probe insertions. If a channel’s signal to noise ratio (SNR), is above threshold (15 a.u., inserts to the right of MUA plots), the corresponding receptive fields (RF) are represented (right plots) with the identical color-code. Note the retinotopic coverage obtained with tangential insertions capturing almost the entire extent of the screen. **C**. Four different sequential stimuli were used: 6 top wedges, 6 bottom wedges, 7 horizontal bars, and 14 vertical bars (top). Wedge exposure is illustrated in time (bottom left) showing the corresponding sequential RF entrainment (bottom right). **D**. Estimate of pairwise phase consistency (PPC) of broad and narrow waveform neurons in 6 different mice, for 3 different repeat frequencies (n= 1115 BW neurons, n = 443 NW neurons, pooled within each of the 6 mice, two ways ANOVA, p_stim = 5.1 10^−3^ for NW, p_stim = 2.0 10^−3^ for BW, with post-hoc repeated measures. Graph shows mean and standard error of the mean (SEM) calculated from the illustrated dots which represent average neuronal responses per mouse. **E**. The underlying working hypothesis is that spatially more complex sequential stimuli can entrain neurons at higher frequency compared to synchronous full-field flickers.

To address whether sequential stimulation can evoke high frequency activity in the visual cortex, we monitored neuronal activity in the primary visual area (V1) of awake head-fixed mice using Neuropixels probes^9^. We accustomed the animal to peacefully sit and watch different visual stimulations. Neuropixels probes inserted in two different angles (Fig. 1A) precisely monitored visual evoked neuronal activity throughout the cortex (Fig. 1B). As routinely done in laboratories studying vision, and as detailed in previous publications^10,11^, great care is taken first to well align the screen position to the measured retinotopy of the current insertion, in order to maximize retinotopic coverage (Fig. 1B). Four types of sequential visual stimulations were presented in a predetermined order, at three different screen frequencies (refresh frame rates = 144/180/240 Hz) for 2 s long trials, in 50 consecutive trials per stimulus type and screen frequency (Fig. 1C). Consequently, a given stimulus having N sequentially presented sections has a duration of N*(1000/screen frequency) ms (Fig. 1C). Given the three different screen frequencies and the 3 different sequential section numbers (6, 7, 14) there were 9 different “repeat frequencies” ranging from 10.28 to 40 Hz, distributed among 12 different sequential stimulations (Ext. Dat. 3). To examine basic response properties, we first measured evoked responses to the different repeat frequencies of the sequential stimuli.

To do so, we extracted the single-unit activity from the different recordings and distinguished them based on their waveform shape into either narrow waveform (NW) putative fast-spiking inhibitory neurons or broad waveform (BW) putative regular spiking excitatory neurons (Ext. Dat. Fig. 1, Ext. Dat. Fig. 2A). In line with previous publications^6^, we report a significant decrease in firing rate with increasing repeat frequencies in all conditions within each individual recording (Ext. Dat. Fig. 2B) and visual stimulus type (Ext. Dat. Fig. 2B, low); similar tendencies were observed in the averages of all recordings of all animals (Ext. Dat. Fig. 2C). First, we observed the 1/f attenuation of neuronal firing in both narrow and broad waveform neurons (Ext. Dat. Fig. 2A-B) in response to the different repeat frequencies of our sequential visual stimulations (regardless of their shape). Second, we quantified the pairwise phase consistency (PPC) of V1 neurons for each repeat frequency to each sequential stimulation and found a significant decrease with increasing repeat frequencies of our sequential stimulations (Fig. 1D, for repeat frequencies of 10/24/40 Hz, BW mean PPC = 0.0879 ± 0.0085/ 0.0251 ± 0.0030/ 0.0012 ± 0.00045 and NW mean PPC = 0.0870 ± 0.0095/ 0.0138 ± 0.00339/ 0.0055 ± 0.0011; two ways ANOVA followed by post-how repeated measure with Bonferroni correction p_values = 0.008/0.0023/0.162 for paired comparison between BW and 0.0149/0.018/0.24 between NW). The details for the twelve different stimulation conditions are reported in the supplementary materials (Ext. Dat. Fig. 3). We hypothesized that the spatially complex sequential structure of our stimulus would evoke an artificial synthetic neuronal wavefront (Fig. 1E).

To answer whether sequential visual stimulation can produce high frequency events, we analyzed the local field potential (LFP) evoked by the different stimulations (Fig. 2A). First, we confirmed the consistent presence of our stimulus repeat frequency in all conditions (Fig. 2A & C for vertical bars, repeat frequency = 10 Hz, Ext. Dat. Fig. 4D & F bottom wedges, repeat frequency = 24 Hz). Then, we quantified in each recording the evoked power spectrum in each channel (Fig. 2B), in order to extract the Z-score of the evoked spectrum (Fig. 2C) for the different sequential stimulations (along with other standard visual stimulations, Fig. 2D, Ext. Dat. Fig. 4). Most of the sequential stimulations entrained a high frequency activity in V1 in defined locations along the recorded retinotopy (Fig. 2C-D). These evoked high frequency activities (between 100-190 Hz) were often present over larger areas in our tangential recordings compared to vertical recordings (Fig. 2D). Its presence confirms that the spatial sequential visual stimulation produced spatially determined entrainment in the cortex. Interestingly, it seemed that lower frequency repeats in our sequential stimulations had a better capacity to entrain such high frequency activity (Fig. 2D) which is likely related to the longer exposure time resulting from the lower exposure time e.g. 4.16 ms for 240 Hz screen frame rate vs. 6.94 ms at 144 Hz (Fig. 1C) and the higher PPC of the lower repeat frequency (Fig. 1D). In order to verify that this high frequency power increase was specific to sequential stimulations, we compared the evoked high frequency activity in the very same recordings to two other classical visual stimulations: a moving bar (white bar on black background) and an alternating full-field stimulus (lower frequency, i.e. 1 to 10 Hz, from Chirp stimulus). The alternating full-field stimulus from 1 to 10 Hz was already sufficient to evoke gamma responses up to 60 Hz (Ext. Dat. Fig. 4C), which are also evoked during most other types of visual stimulations, but incapable of producing this increase of power in the high frequency range.

**Figure 2:**
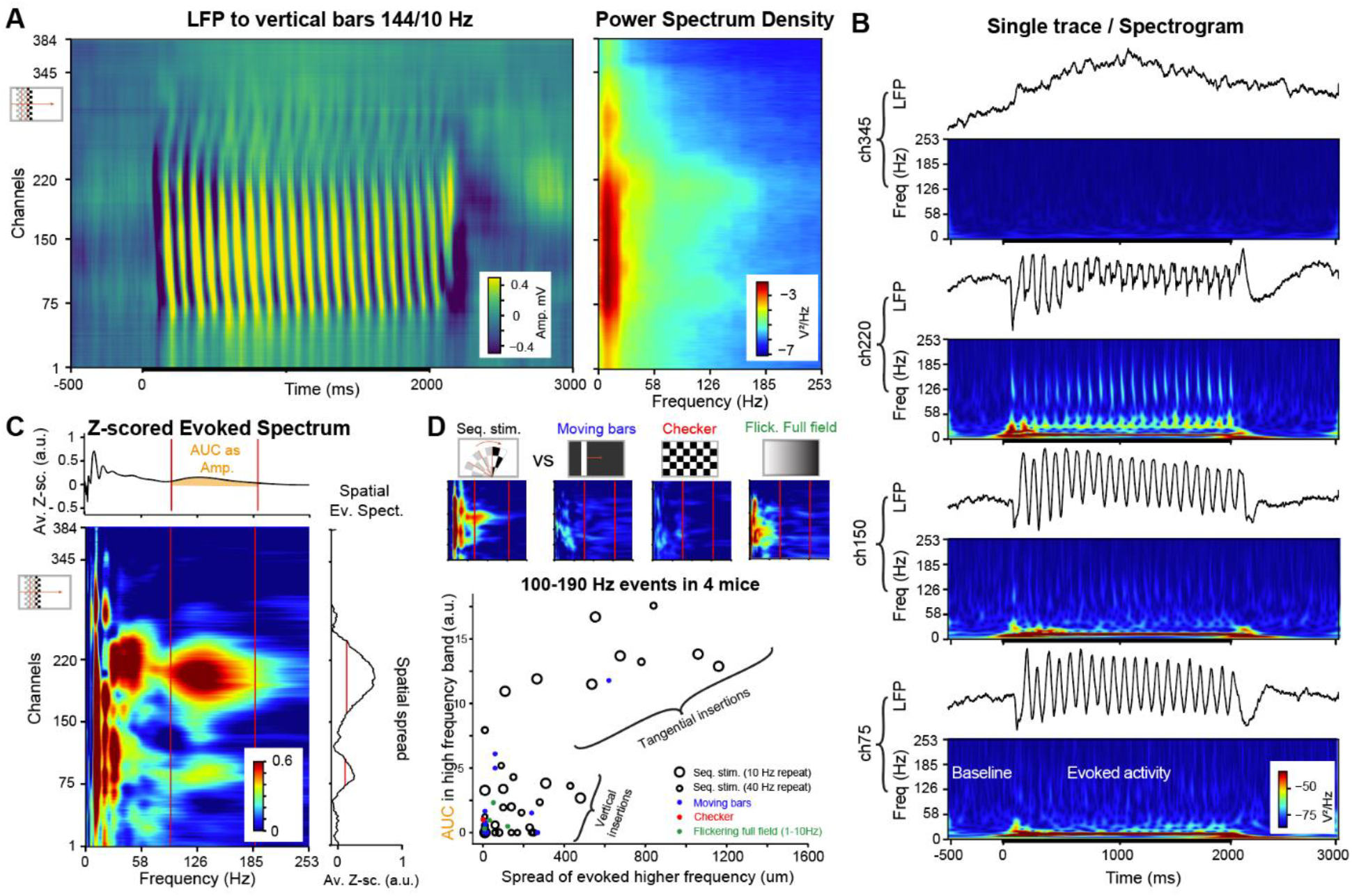
Sequential stimulations enhance high frequency (100 to 190 Hz) power in V1. **A**. Stereotypical evoked LFP averaged across 50 trials of the 2s sequential exposure of vertical bars (left), and the corresponding power spectral density (PSD; right), for the 144 Hz frequency, corresponding to the 10.3 Hz repeat frequency. **B**. Visually evoked LFP (top) and corresponding spectrum (heatmap, bottom), for four different channels along the probe. **C**. Z-scored evoked spectrum based on stable evoked activity, with the corresponding averages in both top and right inserts. These are used to quantify the strength of the evoked high frequency events by the area under the curve (AUC) and the spatial spread of the evoked high frequency events. **D**. Several sequential stimuli and certain moving bars were capable of entraining higher frequency activity in V1 over a large area.

In conclusion, these results from a limited dataset highlight the possibilities for neuronal entrainment beyond full-field flickers. Our results from both single-units and LFPs confirm that lower sequential repeat frequencies entrain stronger responses, hence corroborating the idea that the visual system acts as a low-pass filter. This power loss has been hypothesized to stem from multiple origins, such as synaptic and ion channel noise^12^, active membrane current^13^, neuronal morphology^14^, extracellular space^15^, or even putatively because neuronal capacitance is non-ideal^16^.

Other studies have shown that artificial induction or enhancement of specific brain oscillations can improve cognition. For example, the use of transcranial electric stimulation at 1 Hz during sleep has been shown to enhance memory in healthy humans by increasing slow oscillations and slow spindle activity^17^. 40 Hz full-field flickers improved memory through entrainment of gamma oscillations^2^, although others have challenged such effects^5,6^. Targeting gamma could pose a problem because gamma oscillations are a family of oscillations which have been implicated in many different functions. For example, in mice, broadband and narrowband gamma seem to be two independent oscillations sometimes working in an opposing way^18^: Narrowband gamma are enhanced during running^19^, while broadband gamma are decreased/weakened during running^20^. Similarly, subtle differences have been observed in humans between sensory evoked and cognition induced gamma, hypothesized to underlie the construction of coherent information representation in possibly top-down manners^21^.

In humans, full-field visual stimulation can entrain steady-state visual evoked potentials (SSVEP) up to approximately 90 Hz^22^. This is likely due to the higher capability of human retinal ganglion cells to respond at such high frequency^23^, compared to mice^6^. Our results intend to give a perspective toward more diverse ways to artificially entrain the brain toward therapeutic effects. This approach is bypassing the listed limits of full-field single-frequency repeats, hence being able to induce oscillations at higher frequency. Interestingly, high frequency sequential stimulation at 180 and 240 Hz failed to increase the high frequency power as much as stimulation at 144 Hz (Fig. 2D). This could be due to secondary representations such as orientation tuning and motion coherence, as suggested to occur for gamma oscillations^24^, which should be studied further. Alternatively, this localized high frequency power increase may correspond to the superposition of the two existing wavefronts between the interconnected primary and secondary visual areas, as the stimulus is sequentially exposed. The high PPC values for the 144 Hz stimulus suggest that a certain level of neuronal entrainment occurs in V1, which may feed this high frequency power through secondary cortico-cortical connections. Overall, we cannot tell which exact mechanism(s) are producing this enhancement of high frequency power, as it is variable from one stimulus and one animal to the next. We nevertheless observed a significantly large area of tissue in most of the animals exhibiting an increase of power in high frequency in the visual cortex.

Finally, we propose to extend such sequential approaches to other sensory modalities. Gamma entrainment has already been reported to have different putative therapeutic effects with tactile^25^ and visuo-audio pairing^26,27^. There is a report of pure auditory gamma entrainment using audio stimulation, by multiplexing a 5 kHz carrier into a 40 Hz intensity modulation^28^. Therefore, exploiting the full range retinotopy, tonotopy and somatotopy of sensory stimulation by sequentially activating neighboring receptive fields could become a novel approach to exploit a broader diversity of non-invasive sensory stimulation approaches.

## MATERIAL AND METHODS

### Experimental preparation, animals, Neuropixels probe insertion

All experiments were performed with approval from the local authorities (Landesamt für Gesundheit und Soziales Berlin - G 0142/18). Great care was taken to minimize the number of animals used, which were all recorded in awake head-fixed conditions. The local breeding facility (Charité-Forschungseinrichtung für Experimentelle Medizin) provided the 6 adult C57BL/6J mice, aged 3 to 6 months. A head-post was implanted 10 days prior to recording, under anesthesia (isoflurane, 2.5% in oxygen Cp-Pharma G227L19A), and complemented with painkillers (metamizole in drinking water, renewed daily) prior to and after implantation (−1+3 days). The head post was fixed with dental cement, which was also used to form a bigger circular crown around the planned insertion to retain PBS during recording. The Neuropixels external ground was bound to an Ag/Cl wire fixed directly within this crown to be used as grounding. Habituation was done with daily sessions on the set-up from 30 s to 2 hours gradually increased over 5-7 days. Mice were exposed to sparse noise and moving bars a few days prior to the recording day for them to habituate and peacefully stay in head-fixed condition.

On the recording day, isoflurane was first applied to induce a light anesthesia and perform a small craniotomy, either above V1 for vertical insertions, or above the midline areas for tangential insertions, meanwhile adding a ground cable in the crown. The animal could then recover for at least two hours before being placed on the set-up. Stereotactic coordinates were measured from lambda, in the antero-posterior (AP), the medio-lateral (ML), and the dorso-ventral (DV) axes. Vertical insertions were done with a 30°/30° angle from vertical and ML planes, placed in the brain 0 to 0.9 mm AP, +3 to +4 mm ML (no DV as pseudo vertical). Tangential insertions were done perpendicular to the midline plane with 15-20° from the azimuthal plane; they enter from the right brain side directly above the retrosplenial cortex: 0 to 0.9 mm AP, −0.5 to +0.5 mm ML and 0 to - 0.2 mm DV from lambda (cf. Fig. 1A) as recently published^29^. For both insertions, the probe was placed 4 mm within the brain and then withdrawn 50 µm to release mechanical pressure. The probe was left to settle for ~10-20 minutes before visual stimulations were presented.

Once the recording was finished, the mice were sacrificed with an excess of isoflurane (>4%). Cardiac perfusion started with phosphate buffer saline solution (PBS) and was followed with 4% paraformaldehyde (PFA) and the brain was sliced and mounted on the next days. 3D reconstruction was done using SHARP-track to relate the probe’s positioning to the Allen Mouse Brain Common Coordinate Framework (Fig. 1A).

### Hardware, software

Neuropixels probe (version 1.0) signals were recorded with a PXI system (National Instrument NI-PXIe-1071) using Open Ephys software (www.open-ephys.org). Ecephys was used to calculate isolation distance and refractory period violations (Id >10, Refractory period violation < 0.05% https://github.com/AllenInstitute/ecephys_spike_sorting). The rest of the data analysis was performed in Python 3 (www.anaconda.com), using either Wilcoxon signed-rank tests with Bonferroni correction when looking into single recording data; where the different neurons are considered as independent (Ext. Dat. Fig. 2). Or, for the rest of the statistics quantifying the effects in the population of mice, two-ways ANOVA, followed by repeated measures with Bonferroni correction was used, assuming independence between mice, therefore pooling all neurons within each mouse (Fig. 1, Ext. Dat. Fig. 2 & 3). “***” indicates a p-value below 0.001, “**”a p-value below 0.01, and “*” a p-value below 0.05.

### Sequential and standard visual stimulation

The visual stimuli were displayed on a 27-inch gaming monitor screen (Samsung Odyssey G7 27), at different refresh rates: 120 Hz for standard stimulation and 144/180/240 Hz for the sequential stimulations. The standard stimulus contained a “sparse noise stimuli” made of either a dark or white square on an opposite background, used to map receptive fields (15 deg size, 2 targets/frames of 100 ms, Fig. 1B, 20 repeats for each position on a grid of 36×22 squares), or moving bars (white bars on black background, 10 deg in width, 12 directions, fixed speed of 90 deg/s), followed by alternating full-field checkerboards. For sequential stimulations, 4 different stimulations were presented: top wedges, bottom wedges, vertical bars, and horizontal bars (Fig. 1C). For all sequential stimulations the screen was split into 6, 7, or 14 different sections. The stimulation consisted of alternating black and white checkerboard sections, which were presented sequentially across all neighboring sections to produce a seemingly moving element over the entire screen at different speeds as the screen frame rate was changed from 144 to 240 Hz (Fig. 1C). Once all sections are sequentially exposed, the stimulus was repeated for a duration of 2 s for 50 trials alternated with 2 s gray screen. Consequently, the repeat frequencies were 10.3/12.9/17.1 Hz for the 14 vertical bars, 20.6/25.7/34.3 Hz for the 7 vertical bars and 24/30/40 Hz for both wedge types, at 144/180/240 Hz respectively (Ext. Dat. Fig. 3 for details).

### LFP analysis

The LFP analysis was performed on the 2.5 kHz signal, starting by applying a band pass Butterworth order 3 filter (Scipy, >0.1Hz, <400Hz), then down sampled at 1250 Hz by averaging neighboring data points. Evoked LFPs were produced by averaging all repeats of the same conditions to further study the 2 s long SSVEP responses observed in the brain. Evoked PSDs were illustrated using the Welch function (Fig. 2A, scipy). The evoked spectrum was extracted on a channel-to-channel basis using a Stockwell function to observe over time the changes of power (https://github.com/claudiodsf/stockwell, Fig. 2B & Ext. Dat. 4). A Z-score of evoked activity was then calculated within each channel between baseline and the SSVEPs period (black thick line Fig. 2B, bottom of each plot from 0 to 2000 ms), and then further quantified as a Z-score evoked spectrum (Fig. 2C). The high frequency band of 100-190 Hz was chosen to quantify the strength of evoked high frequency events. The profile of the Z-score evoked spectrum was then estimated (Fig. 2C) to determine the size and strength of evoked high frequency events.

### Single-unit extraction and analysis

Single-unit sorting was done with Kilosort 2.5 (https://github.com/MouseLand/Kilosort) using MATLAB 2018 (www.mathworks.com). Manual inspection merging and curing clustering was done in Phy2 (https://phy.readthedocs.io/en/latest/). Clusters of high quality were kept after applying quality metrics thresholds (Isolation distance < 10, ISI_violation < 0.05%). Broad waveform (putative excitatory) neurons were distinguished from narrow waveforms (putative inhibitory) neurons based on the shape of the waveforms and their peak-to-trough duration (0.43 ms, Ext. Dat. Fig. 1). For the quantification of the visually driven firing responses of both BW and NW, we first added a threshold on the responsiveness of the neurons. To do so, we calculated a peri-stimulus time histogram synchronized to the onset of the different stimuli. This gave a measurement of the pre-stimulus baseline to the evoked activity which we measured the maximum of. We applied a threshold such that the maximum responses were at least 7.5*SNR of the baseline (Ext. Dat. Fig. 2). In two of our six recordings, behavioral noise artifacts (and possibly bad contact within the crown) were present in the recordings (outside of the analyzed visual stimulations) which strongly impaired the quality of the obtained single units, and hence they were discarded for the firing rate analysis (Ext. Dat. Fig. 2) but kept for the pairwise correlations and LFP analysis.

Pairwise phase consistency (PPC) was estimated by measuring the time/phase difference from a fixed sinusoid signal based on the given stimulus repeat frequency (Ext. Dat. Fig. 3). For each neuron’s spikes, the theta values were estimated as the phase position of the artificial oscillation for the stimulus (Ext. Dat. Fig. 3 A). Pairwise sum was then calculated for each neuron and then averaged within each recording (Ext. Dat. Fig. 3B), and dataset (Ext. Dat. Fig. 3C)^6^.

## Data and code availability

Data and scripts will be provided freely upon reasonable request.

## Acknowledgements

The mouse head schema used in Fig. 1F was adapted from https://doi.org/10.5281/zenodo.3925903. This work was supported by the DFG Emmy-Noether grants KR 4062/4–1, KR 4062/4–2 (J. Kr). J. Ke. and V. H. receive salaries from Nuuron GmbH which develops high-frequency visual stimulation methods. We thank Quentin Perrenoud for his input on the manuscript.

## Authors contribution

J. Ke., V. H, and J.S. conceived and designed the study; J. S. collected the data; J.S., J. Ke., and V. H. analyzed the data; J. Ke. and J.S. wrote the manuscript; C. D. and C.V. advised on the project and revised the manuscript, J. Kr., and D. S. provided funding and material support.

## SUPPLEMENTARY MATERIAL

**Extended Data Figure 1:**
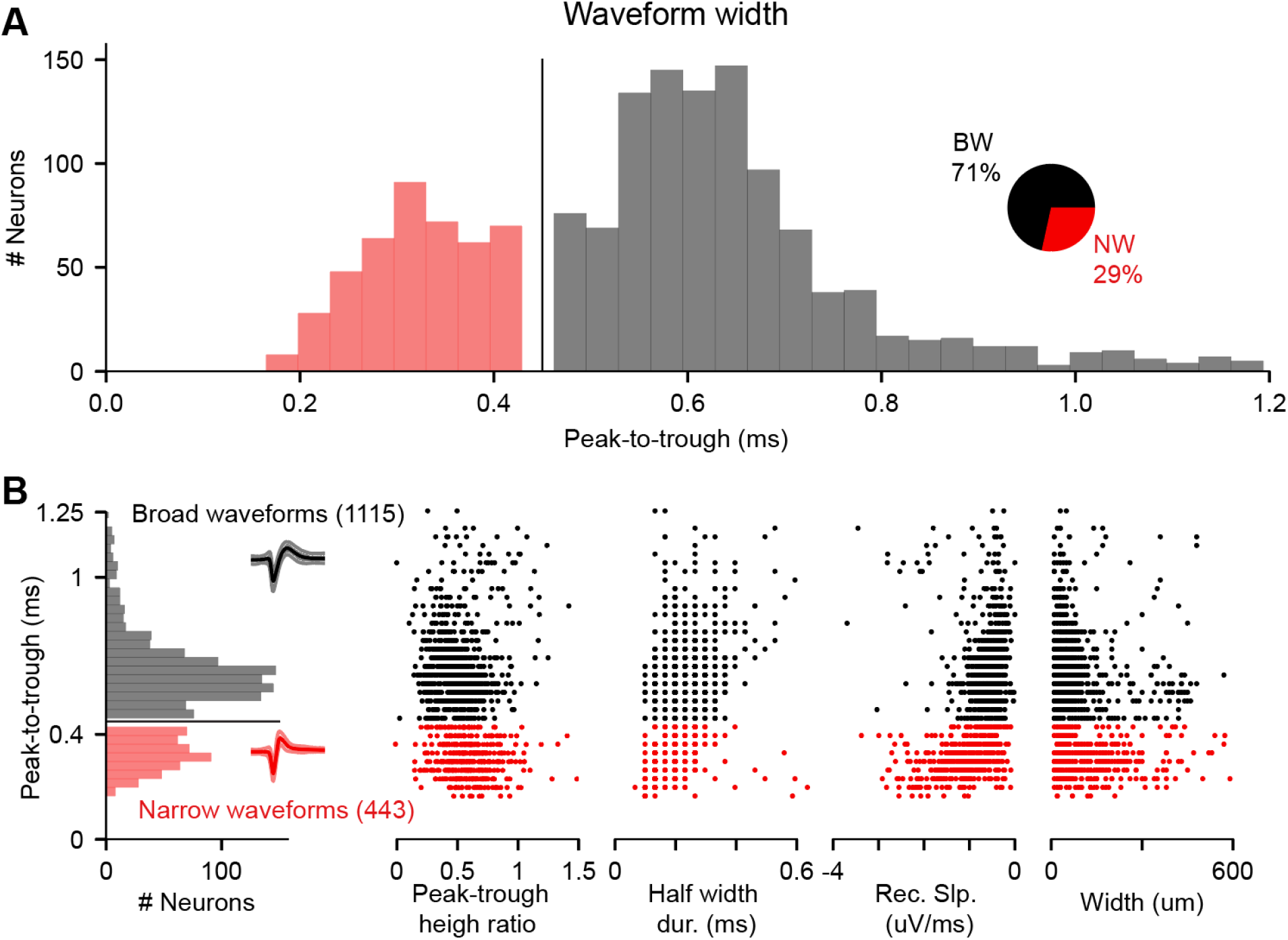
Separation of single-unit waveforms in broad waveforms (BW, putative excitatory) versus narrow waveforms (NW, putative inhibitory) neurons. **A**. Distribution of the waveform duration of the 1558 single-units. **B**. Simple quantification of single-unit properties: peak-to-trough ratio, half-width duration, recovery slope and spatial width.

**Extended Data Figure 2:**
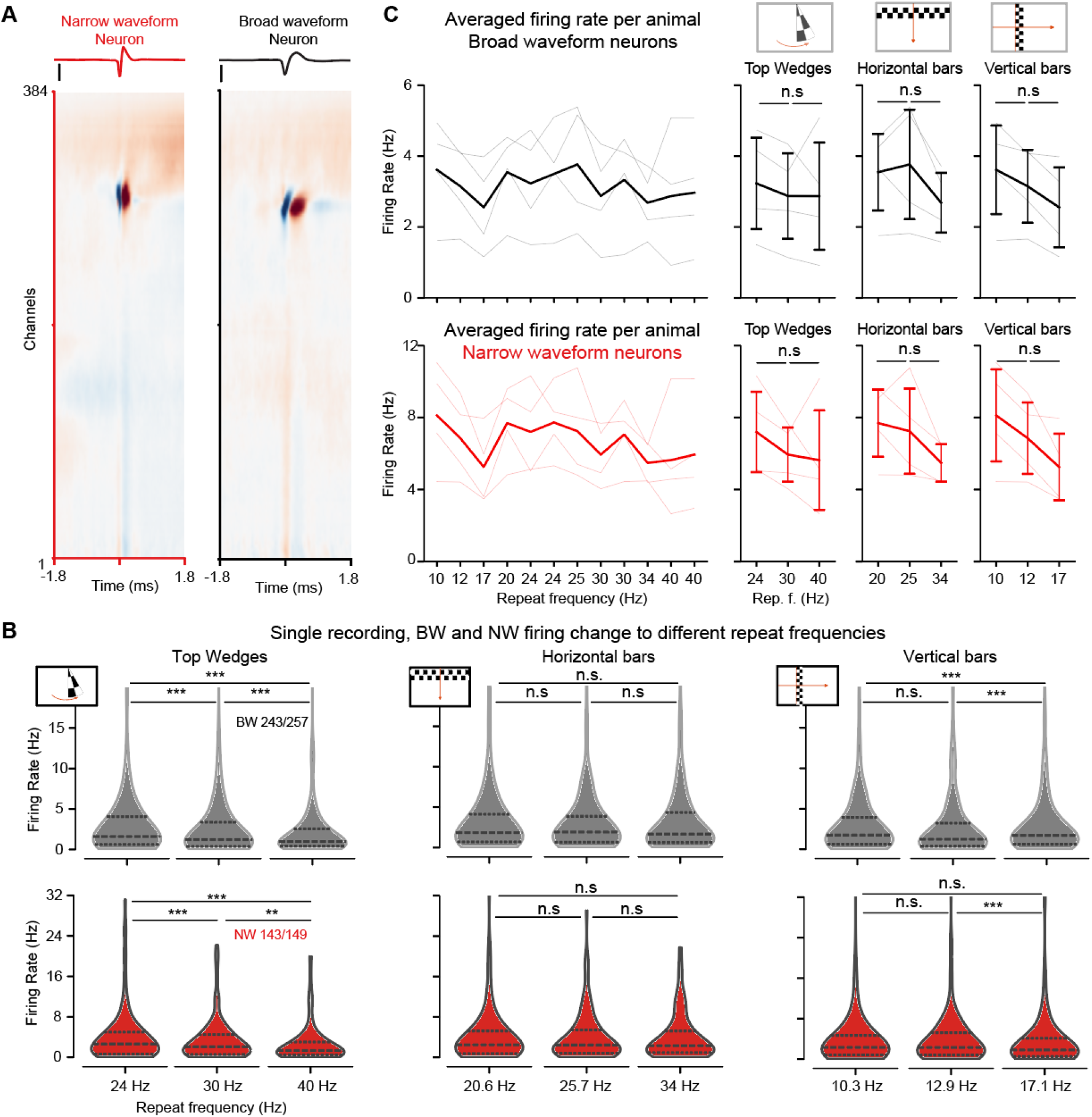
Higher frequency repeats are associated with a reduction of firing of both BW and NW neurons. **A**. Multichannel waveforms of a NW and a BW neuron. **B**. Firing rate changes in a single recording for increasing repeat frequencies in response to top wedges (left), horizontal bars (middle), and vertical bars (right). In both BW neurons (top black), and NW neurons (bottom red), the evoked activity was smaller for higher frequency repeats. Two-sided Wilcoxon signed-rank test (^***^ < 0.001, ^*^ < 0.05), with Bonferroni correction. **C**. Same results in 4 different animals (thin lines) with their respective inter-animal averages (thick lines). (n = 371 NW, n = 951 BW, pooled within each of the n = 4 mice, Two ways ANOVA). Graphs show median, interquartile range (IQR), and range with Seaborn plots, otherwise mean and SEM.

**Extended Data Figure 3:**
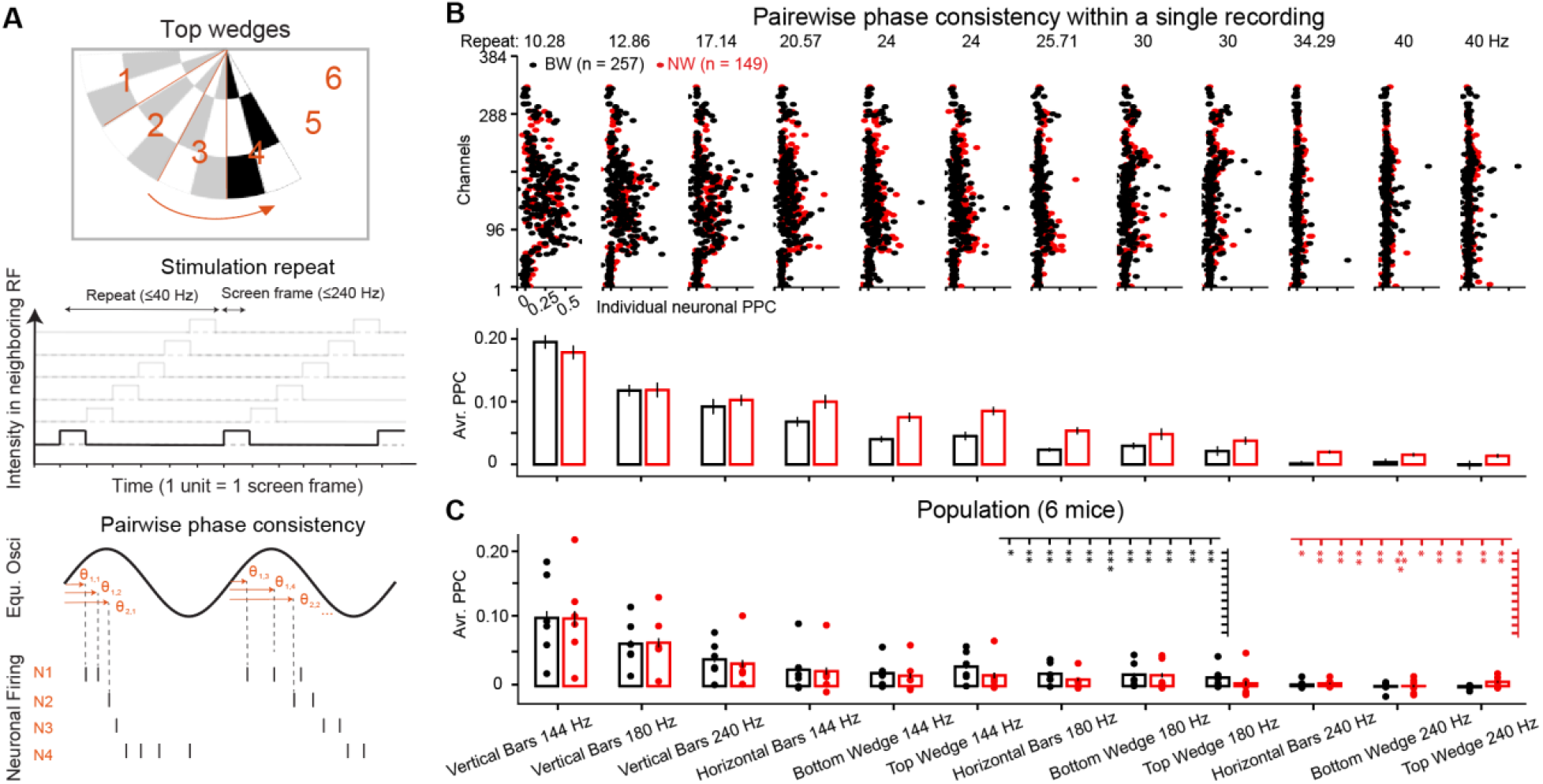
Pairwise phase consistency based on each stimulus repeat frequency. **A**. Schematic of the pairwise phase consistency extraction, one example stimulus (top), the sequential motif resulting from the stimulation (middle), and the equivalent theoretical oscillation upon which the phase of each individual neuron’s spikes is evaluated (bottom). **B**. Channel position of BW and NW PPC (top) and their average population condition (bottom). **C**. Dataset averages of the PPC of BW and NW neurons in V1 along the different repeat frequency content of the different sequential motif combinations, two ways ANOVA, p_stim = 7.3 10^−3^, for BW, p_stim = 8.5 10^−12^, for NW, all neurons. Post hoc repeated measurements with Bonferroni correction report a significant difference (p < 0.05, n = 1115 BW, n = 443 NW, pooled within each of the n = 6 mice) within the vertical bars and toward all others.

**Extended Data Figure 4:**
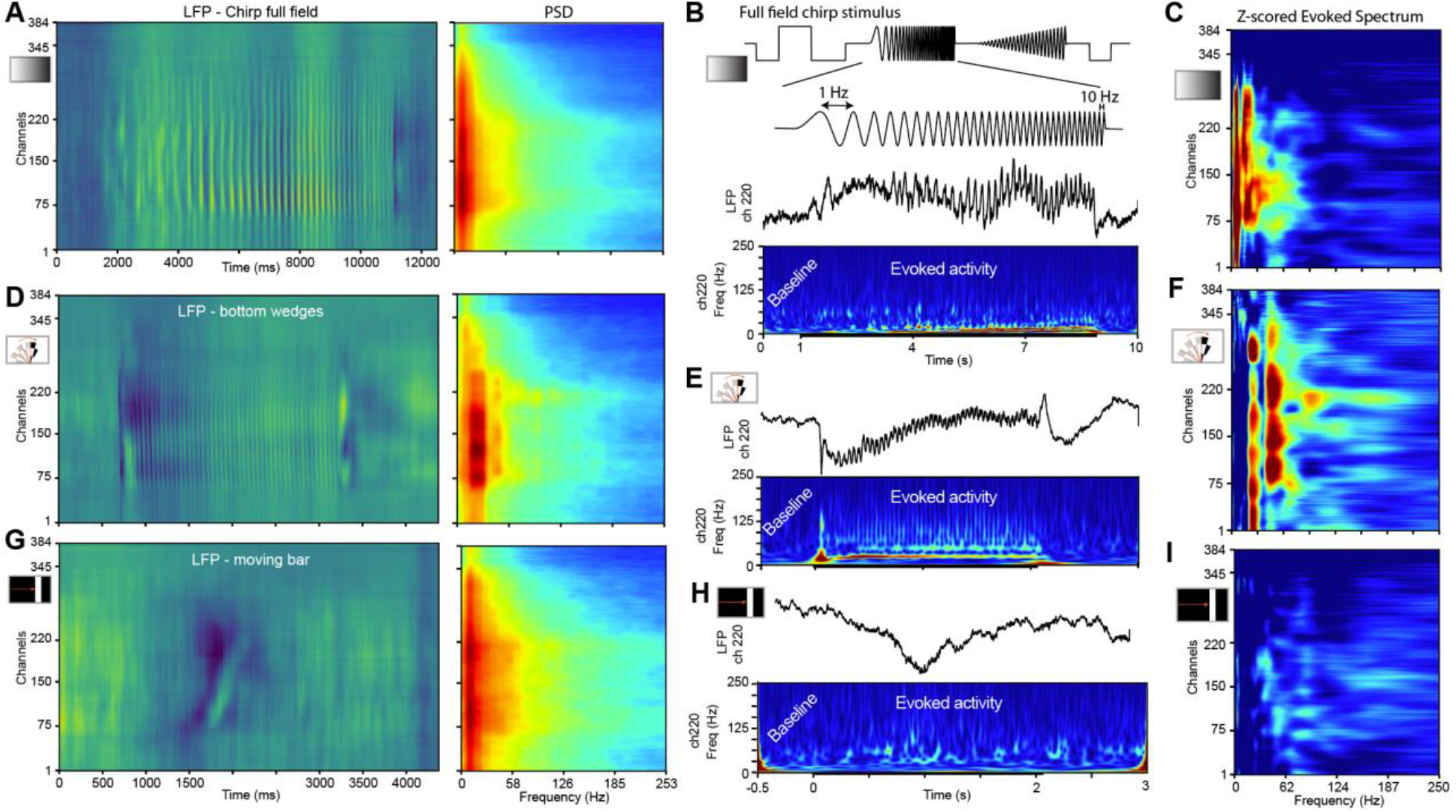
Comparing evoked frequency content of flickering full-field from chirp stimulus and moving bar versus sequential stimulation. **A-C**. Evoked responses from Chirp stimulations. **A**. LFP during the frequency increase of the chirp stimulus (left), PSD of the evoked LFP during a full-field flicker (right). **B**. Full-field chirp stimulus (top) is made of a combination of frequencies and intensity changes, which contain a regular 1 to 10 Hz full-field flicker period (middle top trace), with the corresponding LFP in the most responsive channel (220, lower top trace) and the corresponding spectrogram (lower panel). **C**. Corresponding full-field evoked Z-score (right). **D-F**. Similar plotting as in A-C for sequential bottom wedge stimulus. **D**. LFP, and PSD in response to the bottom wedges stimulus, plotted on similar times as below. **E**. Spectrogram. **F**. Related evoked z-score spectrum. **G-I**. Similar plotting as in **A-C** for moving bar stimulus. **G**. LFP, and PSD. **H**. Spectrogram. **I**. Related evoked z-score spectrum.

## References

1. Iaccarino, H. F. et al. Gamma frequency entrainment attenuates amyloid load and modifies microglia. Nature 540, 230–235 (2016).

2. Adaikkan, C. et al. Gamma Entrainment Binds Higher-Order Brain Regions and Offers Neuroprotection. Neuron 102, 929–943.e8 (2019).

3. Hajós, M. et al. Safety, tolerability, and efficacy estimate of evoked gamma oscillation in mild to moderate Alzheimer’s disease. Front. Neurol. 15, (2024).

4. Zhou, X. et al. 40 Hz light flickering promotes sleep through cortical adenosine signaling. Cell Res 34, 214–231 (2024).

5. Soula, M. et al. Forty-hertz light stimulation does not entrain native gamma oscillations in Alzheimer’s disease model mice. Nat Neurosci 26, 570–578 (2023).

6. Schneider, M., Tzanou, A., Uran, C. & Vinck, M. Cell-type-specific propagation of visual flicker. Cell Reports 42, 112492 (2023).

7. Buzsáki, G., Anastassiou, C. A. & Koch, C. The origin of extracellular fields and currents — EEG, ECoG, LFP and spikes. Nat Rev Neurosci 13, 407–420 (2012).

8. Keil, J. et al. Artificial sharp-wave-ripples to support memory and counter neurodegeneration. Brain Research 1822, 148646 (2024).

9. Jun, J. J. et al. Fully integrated silicon probes for high-density recording of neural activity. Nature 551, 232–236 (2017).

10. Sibille, J., Gehr, C., Teh, K. L. & Kremkow, J. Tangential high-density electrode insertions allow to simultaneously measure neuronal activity across an extended region of the visual field in mouse superior colliculus. Journal of Neuroscience Methods 376, 109622 (2022).

11. Sibille, J. et al. High-density electrode recordings reveal strong and specific connections between retinal ganglion cells and midbrain neurons. Nat Commun 13, 5218 (2022).

12. Pettersen, K. H., Lindén, H., Tetzlaff, T. & Einevoll, G. T. Power Laws from Linear Neuronal Cable Theory: Power Spectral Densities of the Soma Potential, Soma Membrane Current and Single-Neuron Contribution to the EEG. PLOS Computational Biology 10, e1003928 (2014).

13. Reimann, M. W. et al. A Biophysically Detailed Model of Neocortical Local Field Potentials Predicts the Critical Role of Active Membrane Currents. Neuron 79, 375–390 (2013).

14. Pettersen, K. H. & Einevoll, G. T. Amplitude Variability and Extracellular Low-Pass Filtering of Neuronal Spikes. Biophysical Journal 94, 784–802 (2008).

15. Bedard, C., Piette, C., Venance, L. & Destexhe, A. Extracellular and intracellular components of the impedance of neural tissue.

16. Bédard, C. & Destexhe, A. A Modified Cable Formalism for Modeling Neuronal Membranes at High Frequencies. Biophysical Journal 94, 1133–1143 (2008).

17. Marshall, L., Helgadóttir, H., Mölle, M. & Born, J. Boosting slow oscillations during sleep potentiates memory. Nature 444, 610–613 (2006).

18. Saleem, A. B. et al. Subcortical Source and Modulation of the Narrowband Gamma Oscillation in Mouse Visual Cortex. Neuron 93, 315–322 (2017).

19. Vinck, M., Batista-Brito, R., Knoblich, U. & Cardin, J. A. Arousal and Locomotion Make Distinct Contributions to Cortical Activity Patterns and Visual Encoding. Neuron 86, 740–754 (2015).

20. Veit, J., Handy, G., Mossing, D. P., Doiron, B. & Adesnik, H. Cortical VIP neurons locally control the gain but globally control the coherence of gamma band rhythms. Neuron 111, 405–417.e5 (2023).

21. Tallon-Baudry, C. & Bertrand, O. Oscillatory gamma activity in humans and its role in object representation. Trends Cogn Sci 3, 151–162 (1999).

22. Herrmann, C. S. Human EEG responses to 1–100 Hz flicker: resonance phenomena in visual cortex and their potential correlation to cognitive phenomena. Exp Brain Res 137, 346–353 (2001).

23. Schneeweis, D. M. & Schnapf, J. L. Photovoltage of Rods and Cones in the Macaque Retina. Science 268, 1053–1056 (1995).

24. Pellegrini, F., Hawellek, D. J., Pape, A.-A., Hipp, J. F. & Siegel, M. Motion Coherence and Luminance Contrast Interact in Driving Visual Gamma-Band Activity. Cereb Cortex 31, 1622–1631 (2021).

25. Suk, H.-J. et al. Vibrotactile stimulation at gamma frequency mitigates pathology related to neurodegeneration and improves motor function. Front. Aging Neurosci. 15, (2023).

26. Martorell, A. J. et al. Multi-sensory Gamma Stimulation Ameliorates Alzheimer’s-Associated Pathology and Improves Cognition. Cell 177, 256–271.e22 (2019).

27. Murdock, M. H. et al. Multisensory gamma stimulation promotes glymphatic clearance of amyloid. Nature 627, 149–156 (2024).

28. Lahijanian, M., Aghajan, H. & Vahabi, Z. Auditory gamma-band entrainment enhances default mode network connectivity in dementia patients. Sci Rep 14, 13153 (2024).

29. Sibille, J., Gehr, C. & Kremkow, J. Efficient mapping of the thalamocortical monosynaptic connectivity in vivo by tangential insertions of high-density electrodes in the cortex. Proc. Natl. Acad. Sci. U.S.A. 121, e2313048121 (2024).

